# The role of PilU in the surface behaviors of *Pseudomonas aeruginosa*

**DOI:** 10.1101/2024.09.12.612798

**Authors:** Jingchao Zhang, Yan Luo, Yiwu Zong, Shangping Lu, Yi Shi, Fan Jin, Kun Zhao

**Affiliations:** Center for Medical Genetics, Sichuan Provincial People’s Hospital, University of Electronic Science and Technology of China, Chengdu, Sichuan 611731, China; Frontiers Science Center for Synthetic Biology and Key Laboratory of Systems Bioengineering (Ministry of Education), School of Chemical Engineering and Technology, Tianjin University, Tianjin 300072, China; Guangzhou General Institute of Medical Research, Guangdong, Guangzhou 510000, China; School of Life Science and Technology, University of Electronic Science and Technology of China, Chengdu, Sichuan 610054, China; The Sichuan Provincial Key Laboratory for Human Disease Gene Study and The Institute of Laboratory Medicine, Sichuan Provincial People’s Hospital, University of Electronic Science and Technology of China, Chengdu, Sichuan 610054, China; CAS Key Laboratory of Quantitative Engineering Biology, Shenzhen Institute of Synthetic Biology, Shenzhen Institutes of Advanced Technology, Chinese Academy of Sciences, Shenzhen, Guangdong 518055, China; Institute of Fundamental and Frontier Sciences, University of Electronic Science and Technology of China, Chengdu, Sichuan 610054, China

**Keywords:** *Pseudomonas aeruginosa*, Type IV pili, *pilU*, Twitching, Colony expansion

## Abstract

In *Pseudomonas aeruginosa*, the dynamic activity of type IV pilus (TFP) is essential for various bacterial behaviors. While PilU is considered a homolog of the TFP disassembling motor PilT, its specific roles remain unclear. Using pilus visualization and single-cell tracking techniques, we characterized TFP dynamics and surface behaviors in wild-type and Δ*pilU* mutants. We found that Δ*pilU* cells displayed increased TFP numbers but reduced cell movement and delayed microcolony formation. Interestingly, beyond affecting the twitching motility, Δ*pilU* cells formed a thick multilayered colony edge on semi-solid surfaces, slowing colony expansion. Cell-cell collision responses changed from touch-turn dominance in wild-type to touch-upright dominance in Δ*pilU*, affecting colony morphology and expansion. These findings expand our understanding of PilU’s physiological roles and provide potential targets for developing strategies to control *P. aeruginosa* biofilm formation and virulence.

**IMPORTANCE:** The dynamic function of TFP can help cells to search appropriate spatial positions and perceive external environments, and is crucial for a variety of cell behaviors including cell movement, pathogenesis, natural transformation and biofilm formation. Beyond the current picture of PilU in generating high retraction forces through a PilT-dependent manner, this study revealed more roles of PilU in cell surface behaviors including single cell behavior like twitching and multicellular behaviors such as microcolony formation and colony expansion. These findings expand our understanding on the physiological roles of PilU in the surface behaviors of *P. aeruginosa*, and provide us insight in developing TFP-based methods to control the pathogenicity and biofilm formation of *P. aeruginosa*.

## INTRODUCTION

Type IV pilus (TFP) is critical for a variety of bacterial behaviors, including cell movement, biofilm formation, adhesion, DNA uptake, surface sensing, and virulence (1). One well-recognized feature of TFP is that it can drive a special type of surface motion called twitching through the extension-retraction cycles of TFP (2), which has been observed in a broad range of bacteria species (3–7). In addition to its role in the movement that leads to surface exploration, TFP also plays an important role in the transition from reversible to irreversible surface attachment (8, 9). In fact, in *Pseudomonas aeruginosa* (*P. aeruginosa*) and *Caulobacter crescentus* (*C. crescent*), TFP is considered as one key part of surface sensing (10–13). For instance, A. Siryaporn *et al*. found that in *P. aeruginosa*, TFP contraction allows cells to respond to surfaces of varying hardness and promotes the virulence of surface contact (10). Similarly, in *C. crescentus*, C. Ellison *et al*. showed that upon surface contact dynamic TFP activity stopped while fixation adhesins were produced, which then promoted irreversible adhesion (12). Recent studies further revealed a role of TFP-mediated surface sensing in triggering the initiation of the cell cycle of *C. crescentus* (14, 15).

TFP dynamic activity is critical for its function. In gram-negative bacteria, the TFP assembly system is composed of 10 to 18 different proteins, located in the inner membrane, periplasm, and outer membrane (16). The main component of TFP is a protein subunit called major pilin, while minor pilins, although fewer in number, are also crucial for the assembly or specific functions of pilus (17, 18). Biochemical studies have revealed detailed insights into the extension/retraction mechanisms of TFP, including the reverse rotational movement of PilC (pilus inner membrane core protein) driving the extension or retraction of the fiber, as well as the important roles of PilB (pilus assembling protein) and PilT (pilus disassembling protein) in pilus dynamics (19–22). Although PilU is a homology of PilT, its function is not clear yet. This is partially because PilU has been relatively poorly studied compared to PilT and the phenotypes of *pilU* mutants in different organisms are inconsistent (22–28). For instance, in *P. aeruginosa* and *Dichelobacter nodosus* (*D. nodosus*), loss of PilU has been shown to impair twitching motility (23, 24), while in *Neisseria gonorrhoeae*, PilU mutant cells can still twitch and are capable of DNA transformation (25). In *Vibrio cholerae*, Δ*pilU* mutants show a slightly reduced retraction rate (∼1.3-fold) whereas in Δ*pilT* mutants the retraction rate was reduced to ∼50-fold, indicating that PilT is the true retraction motor of TFP (22, 26). This is also true in other species including *P. aeruginosa*, *Acinetobacter baylyi*, and *Acidithiobacillus ferrooxidans* (22, 26–28). It has been suggested that PilU functions as a PilT-dependent retraction ATPase and the coordination between PilT and PilU may be the mechanism for efficient pilin retraction (22, 26, 27). However, the exact coordination of these two motors during this process is still not fully understood. Moreover, beyond its interaction with PilT, more broader impacts of PilU on bacterial surface behaviors are still need to be explored.

The advances in pilin visualization techniques have greatly facilitated the research on TFP. Traditional fluorescence techniques such as immunofluorescence labeling and labeling with large volume fluorescent avidin compounds have been used to visualize TFP (7, 8, 29–31), but these methods cannot get detailed dynamic information of native (intact) TFP associated activities and do not provide temporal resolution. Succinimide dyes combined with exposed primary amines have also been used to visualize the TFP of *P. aeruginosa*. While this approach allows tracking of dynamic elongation and contraction events, the binding of fluorescent dye to pili is not very specific, and the cell body typically will be dyed too with an even stronger fluorescence intensity due to the larger surface area of cell body compared to pili, which increased the difficulties in analyzing TFP (5). Recently, new techniques including a cysteine-substitution-based technique and a label-free interference scattering (iSCAT) microscopy technique have been developed (12, 27), which enables to directly observe TFP in a real-time and in situ manner with both high temporal and high spatial resolutions.

In this study, we aimed to explore the roles of PilU in regulating *P. aeruginosa* surface behaviors through a direct observation of TFP in-situ and in-real time. Toward this goal, we employed the cysteine-substitution-based technique together with bacterial tracking techniques (32), to examining TFP dynamics, single-cell motility, and collective behaviors in wild-type and Δ*pilU* mutant strains. We first characterized the TFP morphology and TFP-driven twitching motility in different mutant strains to show the PilU effect on single cell behaviors, then through monitoring the microcolony formation of different mutants on glass surfaces, as well as colony expansions on semi-solid agar surfaces, the PilU effect on TFP-based cell-cell interactions and the bacterial collective behavior was studied. Our findings suggest that PilU plays multiple roles in regulating *P. aeruginosa* surface behaviors beyond its known function in twitching motility.

## RESULTS

### *pilU* affects TFP number and locations on a cell surface but not TFP extension and retraction speed

To reveal the role of *pilU* in *P. aeruginosa*, we first characterized TFP morphology and activities in three strains, PAO1P*_BAD-_pilA*S99C (WT containing pili visualization plasmid), Δ*pilU*P*_BAD-_pilA*S99C, and Δ*pilT*P*_BAD-_pilA*S99C (hereafter, they will be referred as WT^m^, Δ*pilU^m^*, and Δ*pilT^m^*, respectively). Figure 1A shows successful TFP staining examples of all three strains. With pili observable under fluorescence microscopy, the distribution, number, length and retraction/extension speed of TFP were measured, and the results are shown in Figure 1B-E. TFP were observed to appear at cell poles and/or cell body part. By classifying the distribution of TFP on cell surfaces into three groups, which are poles-only (including one pole and two poles), body-only and poles and body, we found that in all three strains, pili mainly appeared at poles-only, with a percentage of 72 ± 8% in WT^m^, 84 ± 12% in Δ*pilU*^m^ and 97 ± 8% in Δ*pilT*^m^ (Figure 1B). By further differentiating whether pili are located at one-pole or two-poles among these poles-only events, we found that compared with WT^m^ that has a percentage of 16 ± 4% for two-poles event, Δ*pilU*^m^ displayed a percentage of 31 ± 7%, almost as two times as that of WT^m^, while Δ*pilT*^m^ showed a reduced percentage to be 8 ± 2% (Figure 1B), suggesting different roles of PilU and PilT in the control of TFP distribution along cell bodies. Figure 1C shows the measured histograms of TFP numbers per cell. Compared with WT^m^, both Δ*pilU*^m^ and Δ*pilT*^m^ showed an increased percentage of cells with more TFP appeared (Figure 1C), indicating that the absence of PilU/PilT can lead to hyper-piliation of cells. But Δ*pilT*^m^ cells displayed a larger range of TFP length in 0.5 ∼ 10 μm (Figure 1D), compared with that of WT^m^ and Δ*pilU*^m^ in 0.5 ∼ 4 μm, suggesting that PilT is more important than PilU for controlling TFP length.

**Fig. 1.**
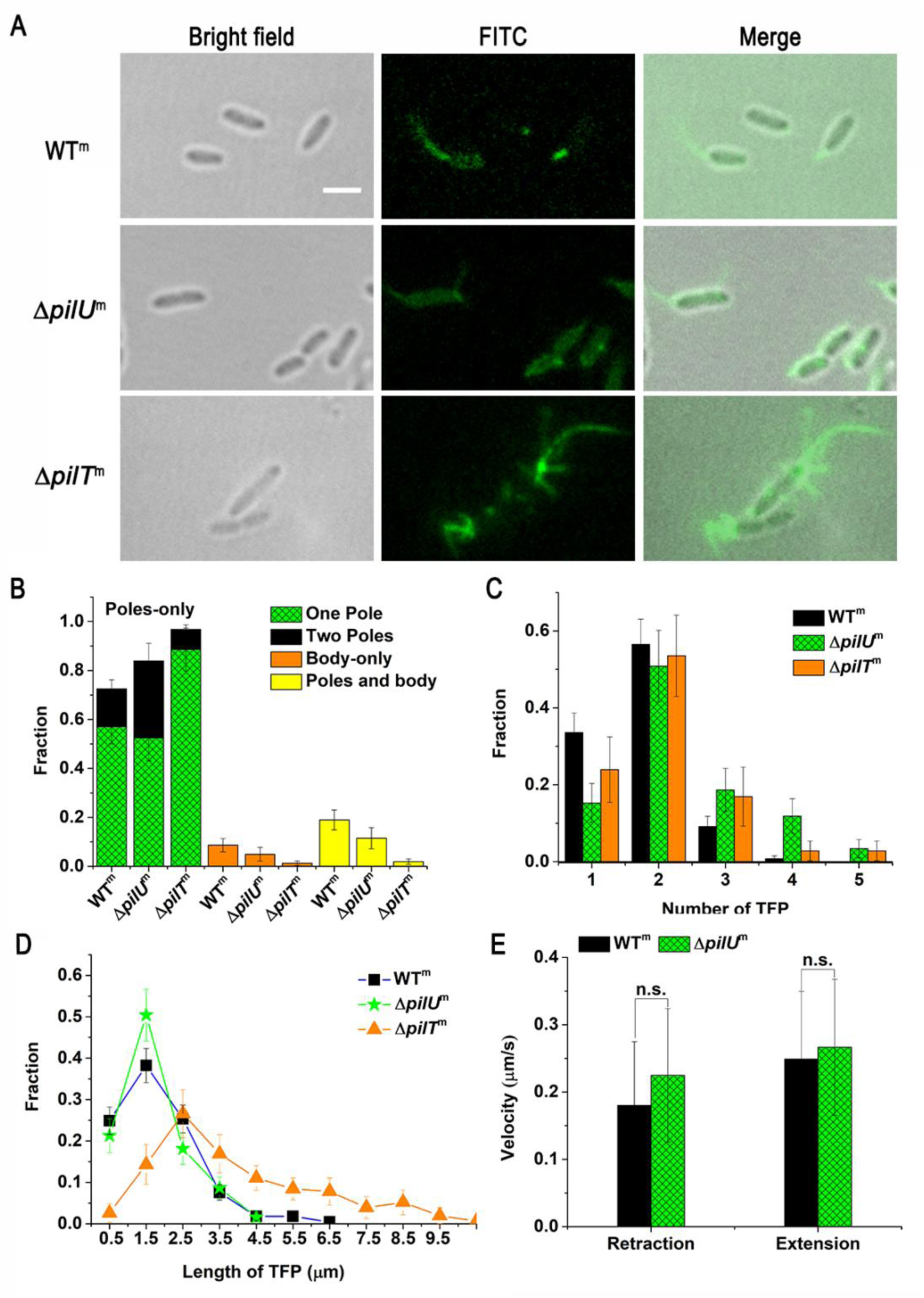
Characterization of TFP morphology of WT^m^, Δ*pilU*^m^, Δ*pilT*^m^. (A) Fluorescence images of stained TFP; (B) The distribution of TFP locations on cell surfaces. *N* (WT^m^) = 116, *N* (Δ*pilU*^m^) = 61, *N* (Δ*pilT*^m^) = 157; (C)The distribution of TFP number per cell. *N* (WT^m^) = 131, *N* (Δ*pilU*^m^) = 59, *N* (Δ*pilT*^m^) = 71; (D) The distribution of TFP length. *N* (WT^m^) = 225, *N* (Δ*pilU*^m^) = 127, *N* (Δ*pilT*^m^) = 131; (E) The extension and retraction velocity, *N* (WT^m^) = 73, *N* (Δ*pilU*^m^) = 28. Error bars show standard deviations. Statistical significances were measured using a two-sample Student’s *t*-test. n.s., not significant; \**p* significant at *p* < 0.05. Scale bars, 2 μm.

Besides morphology, TFP dynamic activity (i.e., extension and retraction of TFP) was also evaluated. In Δ*pilT*^m^, we didn’t observe TFP retraction events, consistent with literature results (27). In contrast, Δ*pilU*^m^ cells still exhibited extension-retraction cycles with an extension speed of 0.26 ± 0.10 μm/s and a retraction speed of 0.22 ± 0.099 μm/s. These speeds showed no statistically significant difference from the corresponding values observed in WT^m^ (0.25 ± 0.10 μm/s for extension and 0.18 ± 0.094 μm/s for retraction) (Figure 1E). This suggests that the absence of *pilU* does not significantly affect the speed of TFP extension and retraction.

### Motile Δ*pilU*^m^ cells displayed a reduced twitching activity but a more directional-persistent surface motion in comparison with WT^m^

Since TFP in *P. aeruginosa* is critical for cell surface movement, we next studied the effects of PilU and PilT on the TFP-driven twitching motility, by analyzing a time series of snapshots that recorded the bacterial surface behavior over a certain time period (see examples in Movie S1-S3). Figure 2A are examples showing the obtained bacterial trajectories for WT^m^, Δ*pilU*^m^ and Δ*pilT*^m^. WT^m^ cells displayed a variety of trajectory patterns as expected. By contrast, Δ*pilT*^m^ cells showed only dot-like short traces. Note that such short traces are artifacts due to the cell elongation through cell growth as their TFP didn’t retract and cells thus didn’t twitch. Δ*pilU*^m^, on the other hand, showed both long straight trajectories, which were definitely generated by cell movement, and short traces. As Δ*pilT*^m^ didn’t twitch, we focus on the comparison between WT^m^ and Δ*pilU*^m^. Although both WT^m^ and Δ*pilU*^m^ cells can twitch, the proportion of motile cells is higher in WT^m^ (79 ± 16%) than in Δ*pilU*^m^ (46 ± 14%) (Figure S1). The trajectories of motile cells, can be quantitatively characterized by mean square displacements (MSDs), which measures to what extent a cell motion deviates from a typical random diffusive motion and whose slope in a log-log plot reflects the shape of cell trajectories. Figure 2B shows the measured MSDs of WT^m^ and Δ*pilU*^m^. The MSD of WT^m^ exhibited a fitting slope of 1.02 ± 0.02, suggesting a random motion of WT^m^ cells. By contrast, the MSD of Δ*pilU*^m^ has a fitting slope of 1.21 ± 0.02, indicating a super-diffusive-like motion. To further understand the different surface motion shown by MSDs between WT^m^ and Δ*pilU*^m^, we calculated twitching speed and the fraction of movement time during the total recording time for each cell (Figure 2C and 2D). Overall, the speed distribution of WT^m^ is right shifted compared with that of Δ*pilU*^m^, indicating that the twitching activity of Δ*pilU*^m^ is reduced compared with WT^m^ (Figure 2C).This is also consistent with the measured result of the fraction of movement time (Figure 2D), which is 0.59 ± 0.09 for Δ*pilU*^m^ (i.e., for a tracked on-average motile Δ*pilU*^m^ cell, there is about 59% of the total tracked time during which the cell moved), but is 0.84 ± 0.08 for WT^m^.

**Fig. 2.**
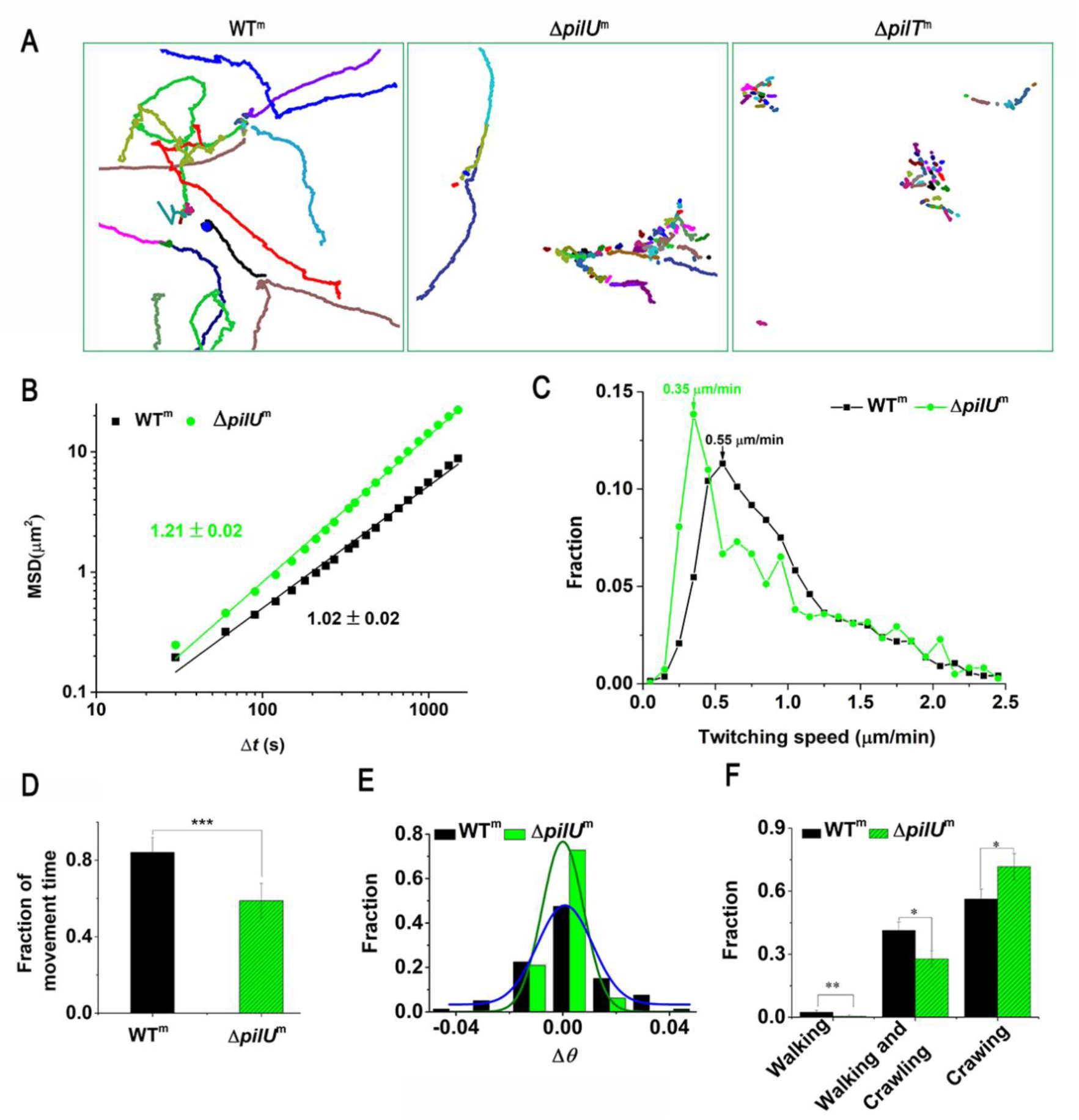
Effects of PilU loss on bacterial surface motility. (A) Examples showing trajectories of *P. aeruginosa* WT^m^, Δ*pilU*^m^ and Δ*pilT*^m^. Different colors represent trajectories of different cells; (B) Mean square displacements; (C) Distributions of twitching speed. *N* (WT^m^) = 7701, *N* (Δ*pilU*^m^) = 2708; (D) Fraction of movement time. *N* (WT^m^) = 49, *N* (Δ*pilU*^m^) = 33; (E) Distributions of Δ*θ*, which is defined as the orientational angle difference of a cell between two consecutive frames. Curves are the Gaussian-Fitting results. *N* (WT^m^) = 81, *N* (Δ*pilU*^m^) = 81; (F) Distributions of twitching modes. *N* (WT^m^) =494, *N* (Δ*pilU*^m^) = 494. Error bars show standard deviations. Statistical significances were measured using a two-sample Student’s *t*-test. \**p* < 0.05, \*\**p* < 0.01, and \*\*\**p* < 0.001.

As the trajectory pattern is not only cell-speed-dependent but also affected by the directional persistence of cell movement, we examined the orientational angle difference of a cell between two consecutive frames, Δ*θ* (Figure 2E). The results showed that the Δ*θ* distribution of Δ*pilU*^m^ is narrower than that of WT^m^. The Δ*θ* of WT^m^ ranged from −0.04 to 0.04 rad while that of Δ*pilU*^m^ ranged from −0.02 to 0.02 rad. We also counted the distribution of the angle between the cell velocity and the x-axis (as a reference axis) (Figure S2), but there was no distinguished difference between the two strains, which may be due to relatively large fluctuations in the velocity calculations.

Besides Δ*θ*, the proportions of different twitching modes of cells were also measured (Figure 2F), as previous studies have shown that different twitching modes have different MSDs (2, 33). The results show that compared with WT^m^, Δ*pilU*^m^ has a higher percentage of crawling-only mode (70% ± 6% for Δ*pilU*^m^ and 56% ± 5% for WT^m^) and a lower percentage of walking-only and walking and crawling modes. Since walking motility often generates a random diffusive motion, low percentages of walking-involved modes in Δ*pilU*^m^ can help cells to maintain their moving direction. Taken together, these results suggest that compared with motile WT^m^ cells that showed a diffusive-type surface motion, motile Δ*pilU*^m^ cells have a reduced twitching activity, but their more confined Δ*θ* and low percentages of walking-involved twitching modes resulted in a more directional-persistent surface motion characterized by a higher slope of MSDs.

### Loss of PilU affects the microcolony formation time through reduced twitching activity and the microcolony morphology by altered cell-cell collision response patterns

Biofilm formation is a classical collective behavior of cells after they attached on surfaces, during which TFP plays an important role (34). The development of a normal biofilm requires not only the physical pilus itself but also its dynamic activity (35). To understand whether and how PilU can affect the biofilm development, we studied the dynamics of microcolony formation of these three strains (Figure 3). Earlier studies have shown that microcolony formation was closely related to bacterial visit distributions on surfaces through Psl-guided surface movement (36), so we first examined the visit frequency map of cells obtained 4h after bacterial inoculation (Figure 3A), which showed how often a pixel was visited by bacteria during the 4 hours. The color scale in Figure 3A ranges from black to blue, representing the number of bacterial visits from 0 to 100. Among the three strains, WT^m^ has a relatively more uniform distribution while Δ*pilT*^m^ has a more localized one and Δ*pilU*^m^ is kind of in between. Quantitatively, the bacterial visit frequency map can be characterized using a power-law distribution (36), and a more negative power-law exponent would indicate a more uniform distribution. The results in Figure 3B show that WT^m^, Δ*pilU*^m^ and Δ*pilT*^m^ has a power law bacterial visit distribution with an exponent of −2.87 ± 0.11, −2.69 ± 0.15 and −2.58 ± 0.11, respectively. Thus, compared to WT^m^, retraction motor mutants Δ*pilU*^m^ and Δ*pilT*^m^ exhibited more hierarchical distribution of bacterial visits (Figure 3B). Next, we measured the microcolony formation time defined as the time period from the inoculation of bacteria to observing the first microcolony in the field of view. Here, a microcolony was defined as a cell aggregate consisting of ≥ 30 cells, the same definition used in literature (33). The results are shown in Figure 3C. The microcolony formation time is 3.5 ± 0.2 h for WT^m^, 4.7 ± 0.9 h for Δ*pilU*^m^ and 5.1 ± 0.8 h for Δ*pilT*^m^. These results indicate that the loss of retraction motors PilU and PilT would slow down the microcolony formation. Particularly, the loss of PilT (Δ*pilT*^m^) was slowed more than the loss of PilU (Δ*pilU*^m^). As the three strains show similar growth curves (Figure S3), such differences in the microcolony formation time are likely caused by their motility differences. To further reveal the role of motility in the microcolony formation, we traced the cell lineages in the course of microcolony formation using the same method as in (36). The results were shown in Figure 3D, where different colors represent different lineages. The microcolony of WT^m^ consisted of cells with about 10 colors (i.e., 10 cell lineages) while the microcolony of Δ*pilT*^m^ consisted of cells with only 2 colors. This is understandable, as WT^m^ cells can twitch and thus can move to join other cells to form microcolony while Δ*pilT*^m^ cells cannot twitch so their microcolonies were formed from one or two cell lineages through cell multiplication. Similarly, Δ*pilU*^m^ cells can twitch but has a reduced twitching activity compared with WT^m^, thus the microcolony of Δ*pilU*^m^ is in between and is composed of cells with 7 colors. These results indicate that cell twitching mobility can enrich the composition of microcolonies, and speed up the microcolony formation.

**Fig. 3.**
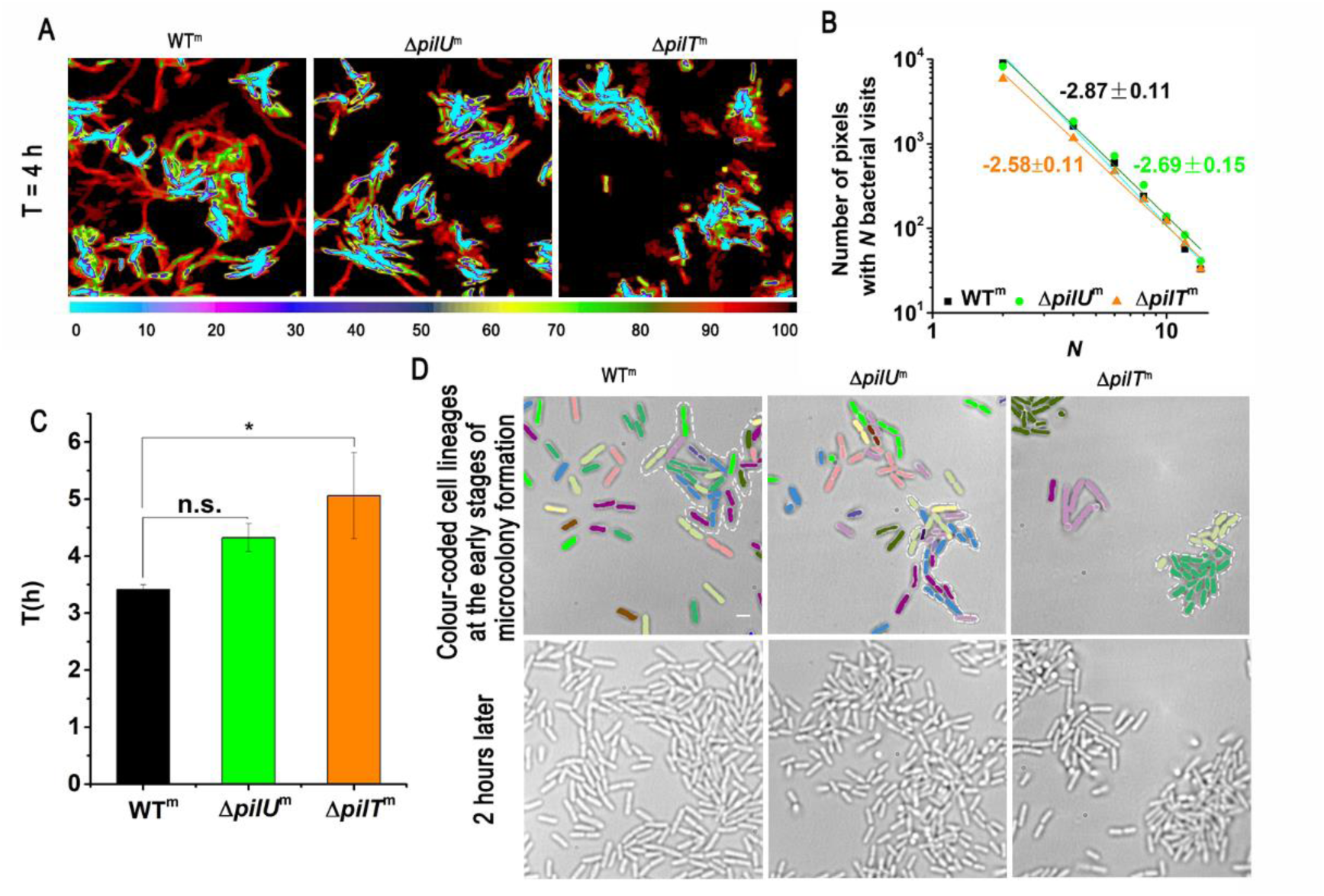
Effects of PilU loss on the formation of microcolonies. (A) Distribution of bacterial visit frequencies measured 4h after inoculation for the three strains; (B) Power law distribution of bacterial visit frequencies shown in (A); (C) The microcolony formation time; (D) The composition of microcolonies formed by the three strains, WT^m^, Δ*pilU*^m^ and Δ*pilT*^m^. Cells in the images of the top row were color-coded, with each color representing a different cell lineage determined from bacterial tracking. In each image, a microcolony was outlined by a white dotted line. The bottom row depicts more developed microcolonies at the same location 2 h later. Scale bars, 2 μm. Error bars show standard deviations. Statistical significances were measured using a two-sample Student’s *t*-test. n.s., not significant; \**p* < 0.05.

The microcolony formation is a multi-cellular process, during which cell-cell interactions are critical for the final formed structures of microcolony. To explore the TFP-mediated cell-cell interactions, we took advantage of the visualization of TFP, and investigated what motile cells would respond when their TFP detected other cells (i.e., their TFP touched with other cells) along their moving trajectories. The results (Figure 4) showed that when cells collided with each other during their movement, generally they exhibited two types of response behavior, touch-turn and touch-upright (see schematic illustrations in Figure 4A). Touch-turn refers to that a bacterial cell changes its orientation and movement direction (but in a plane parallel to the substrate) after the cell collides with another cell. Touch-upright refers to that a cell becomes in stand-up configuration after it contacts with another cell. Figure 4B shows one example for each type of response behavior in WT^m^ and Δ*pilU*^m^ (see movie S4 - S7 for more details). By counting the percentage of cells that displayed touch-turn and touch-upright responses among all observed cell-cell collision events, we found that WT^m^ cells showed more touch-turn responses than touch-upright whereas Δ*pilU*^m^ cells behave just opposite (Figure 4C). Such differences in cell-cell collision responses between WT^m^ and Δ*pilU*^m^ may contribute to the different morphology of microcolonies formed in these two strains (Figure 3D), where in a WT^m^ microcolony most of cells lied down and thus the orientation of cells was aligned to a certain degree due to the crowd packing environment inside the microcolony while in a Δ*pilU*^m^ microcolony quite a few cells stood up so cells orientated more randomly and thus make the microcolony a more-disordered looking.

**Fig. 4.**
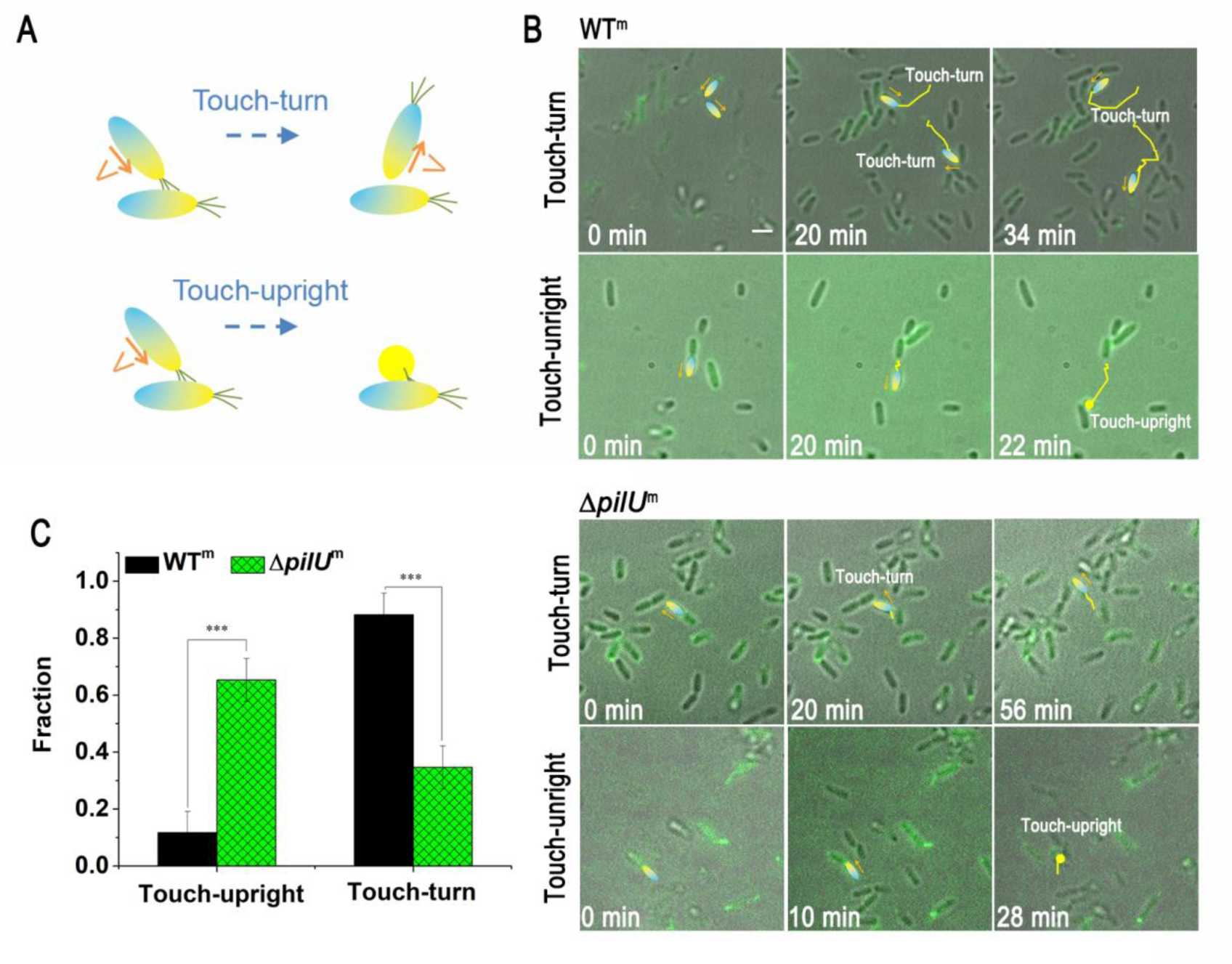
Effects of PilU loss on cell-cell collision responses. (A) The two types of response behavior of cells when they contact with each other. (B) Examples of touch-turn and touch-upright responses in WT^m^ and Δ*pilU*^m^. The interested cells are colored using the same color scheme as in the cartoon in (A), and the yellow lines show their trajectories from 0 min up to the current time point. Orange arrows indicate the moving direction of cells. (C) The measured fraction of touch-turn and touch-upright events among all observed cell-cell collisions. *N* (WT^m^) = 120, *N* (Δ*pilU*^m^) = 51. Scale bars, 5μm. Error bars show standard deviations. Statistical significances were measured using a two-sample Student’s *t*-test. \*\*\**p* < 0.001. Scale bars, 2 μm.

### Beyond twitching, loss of PilU altered colony expansion behavior to form a dense multilayered colony edge when cell population expand on semi-solid surfaces, which consequently slowed down the colony expansion

It has been shown that bacteria behave differently on substrates of varying hardness (37, 38). As PilU is known to play an important role when cells require significant retraction forces (22, 27), cells without PilU may behave differently on semi-solid surfaces. To test this hypothesis, we studied the colony expansion of Δ*pilU*, Δ*pilA*, Δ*pilT* and WT on semi-solid agar surfaces. The results show that the expansion edges of Δ*pilU*, Δ*pilA*, Δ*pilT* and WT displayed different morphologies (Figure 5A). As a control, the expansion edges of WT are rough with relatively large interdigitated convex and concave regions. By contrast, the expansion edges of Δ*pilU*, Δ*pilT* and Δ*pilA* are relatively smooth. A closer examination of bacteria at the front of colony expansion further revealed that Δ*pilT*, Δ*pilA* and WT all have a single-layer of cells at the front of expansion while Δ*pilU* has a multi-layered cellular structure.

**Fig. 5.**
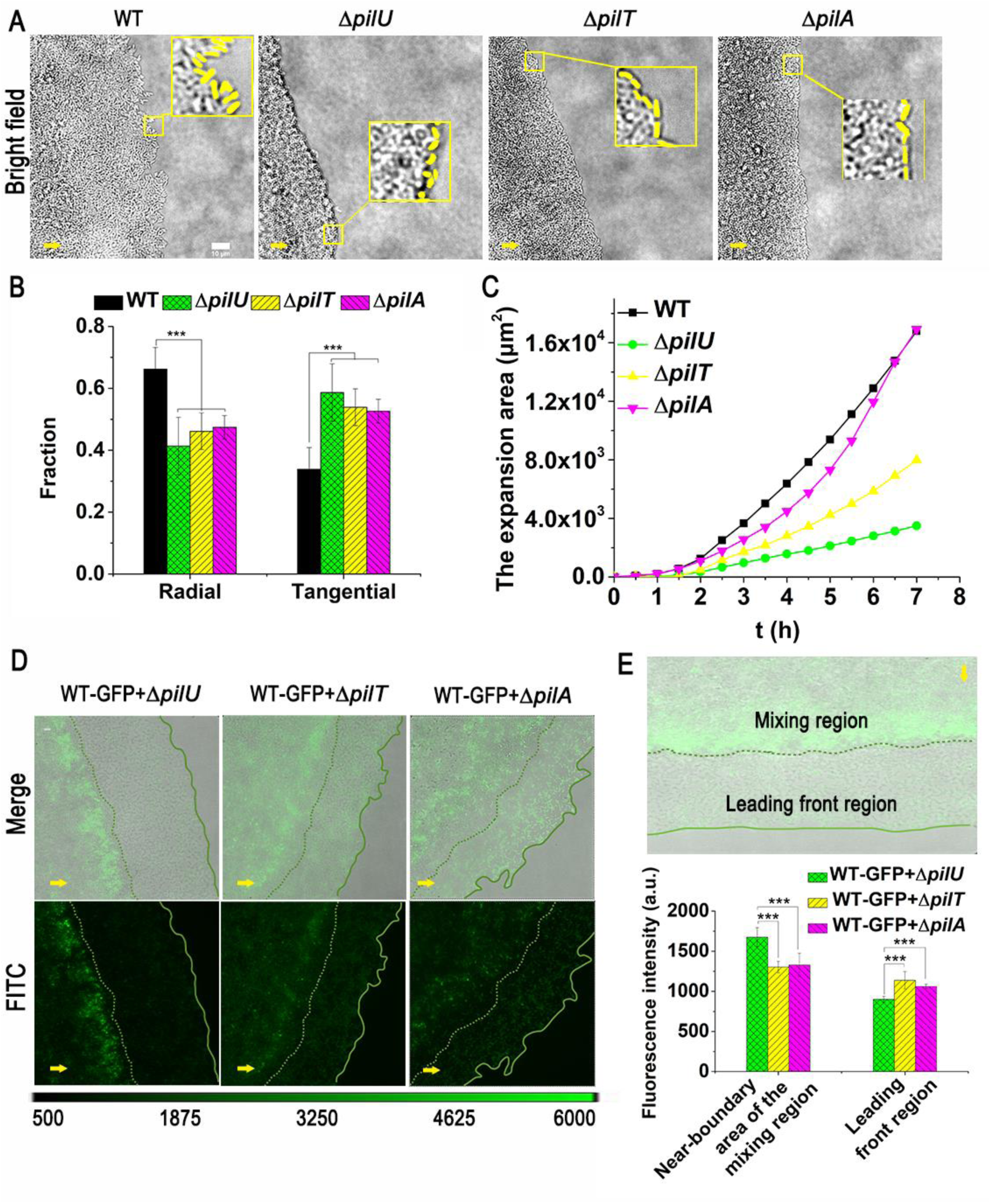
Effects of PilU loss on the colony expansion on semi-solid agar surfaces. (A) Images showing a part of colony expansion edge of *P. aeruginosa* WT, Δ*pilU*, Δ*pilT* and Δ*pilA* after 12 h incubation. Insets are magnified views of specified region at the expansion edge enclosed by yellow squares, in which cells at the edge were marked yellow. The yellow arrow at the bottom left corner of each image indicates the expansion direction. Scale bar: 10 μm; (B) The orientation of cells at the expansion edge. The results were obtained from a total of *N* field of views: *N* = 10 for WT, and the total count number of cells at the edge of community is 601; *N* = 14 for Δ*pilU*, and the total count number of cells is 650; *N* = 13 for Δ*pilT*, and the total count number of cells is 556; *N* = 10 for Δ*pilA*, and the total count number of cells is 619. (C) The expansion area of WT, Δ*pilU*, Δ*pilT* and Δ*pilA*; (D) Distribution of fluorescence signals in the expansion colony of mixed strains. Snapshots were taken after 12 h incubation. Lines are guides to the eye, where the solid line indicates the edge of leading front and the dotted line indicates the boundary between the mixing region and leading front region. The yellow arrow at the bottom left corner of each image indicates the expansion direction. Scale bar: 10 μm; (E) Comparison of fluorescence intensity between the near-boundary area of mixing region and the leading front region. Statistical significances were measured using one-way ANOVE. ** *p* < 0.01, and *** *p* < 0.001.

In addition, the distribution of cell orientations at the colony edge also shows different trend between WT and TFP mutants, as shown in Figure 5B. We can see that compared with WT, TFP mutants including Δ*pilU*, Δ*pilA*, and Δ*pilT* displayed an increase in the percentage of cells that orientated along the tangential direction of the colony edge and a decrease in the percentage of cells that orientated along the radial direction of the colony (i.e. from the colony center towards the periphery). Interestingly, we found that in Δ*pil*A and Δ*pilT*, cells at the colony edge were all in a lie-down style (i.e. ∼ 0% stand-up cells), whereas in Δ*pilU*, more stand-up cells (56 ± 13%) at the colony edge were observed (see examples in the enlarged windows enclosed by yellow rectangles in Figure 5A) (Figure S4). The stand-up configuration of Δ*pilU* is very likely related to the touch-upright responses of cells demonstrated in Figure 4, which may also contribute to the multilayered structure at the colony front in Δ*pilU*. All these differences in the structure and morphology of expansion front among the tested strains, led to different speed of colony expansion with on average PAO1 > Δ*pilA* > Δ*pilT* > Δ*pilU* (Figure 5C).

To further understand the role of TFP-mediated interactions in colony expansion, we mixed WT containing green fluorescent protein (GFP) - expressing plasmids (WT-GFP) with Δ*pilU*, Δ*pilT* and Δ*pilA*, respectively. Each mixture has a ratio of WT-GFP:TFP mutant = 1:5. The results are shown in Figure 5D. We can see that after colony expanded for a certain time, stratification of cells was observed in all tested mixtures.

At the leading front of bacterial colony, the majority of cells were WT-GFP cells and they essentially formed a monolayer. This result suggests that cells with malfunctioned TFP expanded slower than WT, which is in consistent with the results shown in Figure 5C. Behind this WT-dominated leading front region, cells of WT-GFP and TFP mutants were mixed and formed a multilayered structure with high cell density. In this mixing region, the expansion was mainly due to cell growth and multiplication.

Interestingly, WT-GFP cells in the mixing region displayed different distributions among different mixtures. In the WT-GFP + Δ*pilU* mixture, the fluorescence intensity of the mixing region showed a non-uniform distribution with much brighter signals near the boundary, indicating an accumulation of WT-GFP cells at those locations. Whereas in the WT-GFP + Δ*pilT* mixture, the degree of non-uniformity of the fluorescence intensity distribution was less although near-boundary areas were still brighter compared with other areas in the mixing region. By contrast, in the WT-GFP + Δ*pilA* mixture, the fluorescence intensity is more uniformly distributed, indicating that there was no accumulation of WT-GFP cells near the boundary. Quantitatively, by normalizing the fluorescence intensity of the near-boundary area by that of the leading front region, WT-GFP + Δ*pilU* has the highest ratio (∼ 2) (Figure 5E). Considering the observation that a colony of pure Δ*pilU* cells has a multilayered edge (Figure 5A), these results may suggest that Δ*pilU* cells can block the migration of WT cells toward the leading front region through forming a multilayered edge and thus led to the accumulation of WT-GFP cells near the boundary.

We note that the multilayered colony edge phenotype in Δ*pilU* is not due to the affected twitching motility *per se*, as such phenomena was not observed in either WT (normal twitching) or Δ*pilA*/Δ*pilT* (non-twitching). Together, all these results indicate that beyond affecting the twitching motility, PilU also plays a role in the regulation of cell colony expansions on semi-solid surfaces.

## DISCUSSION

Our study reveals multiple roles of PilU in regulating *P. aeruginosa* surface behaviors. We found that PilU influences (1) TFP distribution and quantity, (2) single-cell motility patterns, (3) cell-cell collision responses, and (4) collective behaviors in microcolony formation and colony expansion, which are summarized in Figure 6.

**Fig. 6.**
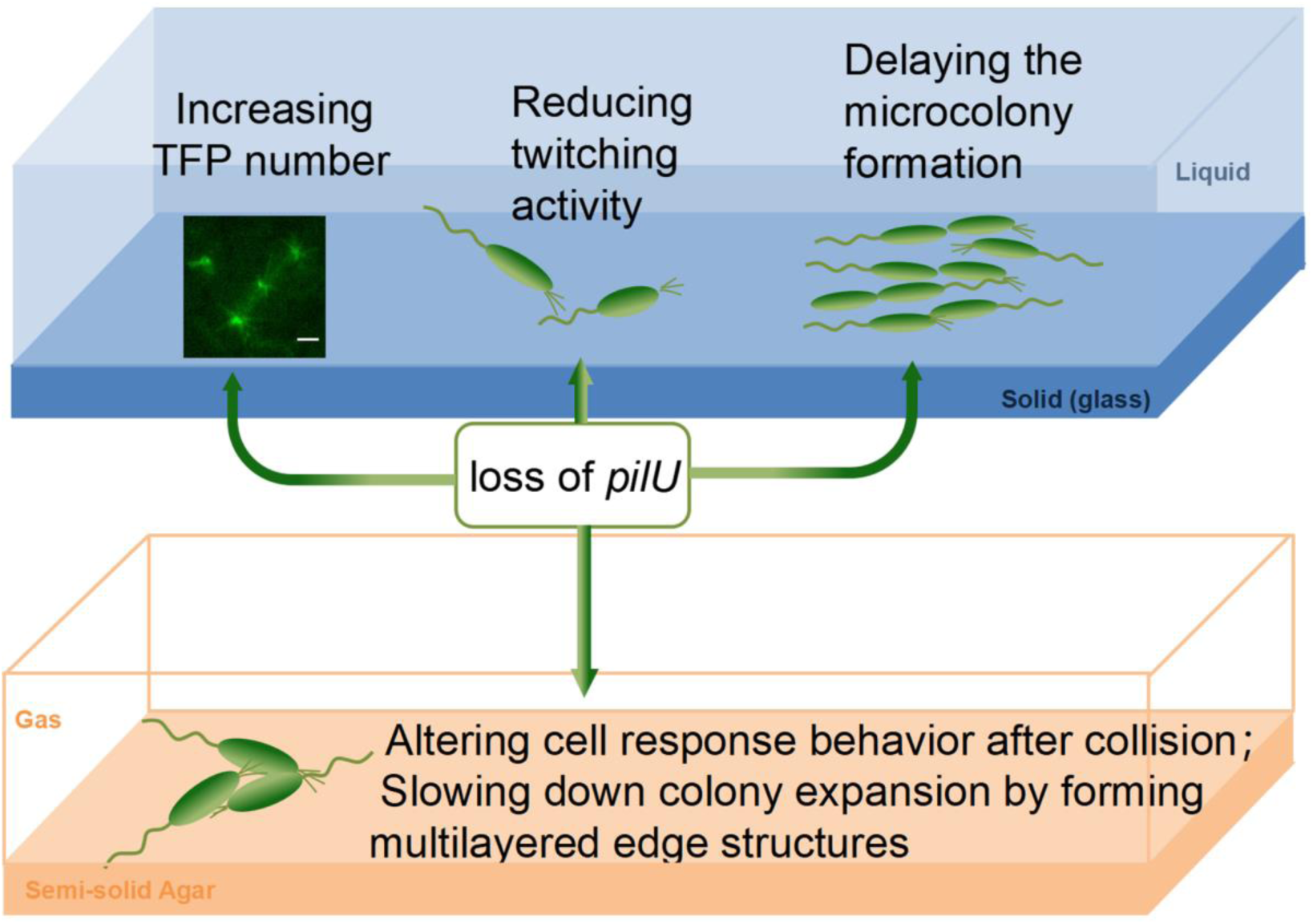
Summary of the effects of *pilU* on the surface behaviors of *P. aeruginosa*.

Firstly, PilU affects cell morphology as loss of PilU increased TFP number on each cell. This is consistent with the hyperpiliated phenotype reported in literature (23, 39). Compared with PilT, our results revealed that PilU played an important role in affecting the quantity of TFP, while PilT is more important in affecting the length of TFP. Consistent with the different impacts on cell morphology between PilU and PilT, they also resulted in different phenotypes with different cell twitching activities, where Δ*pilT*^m^ cannot twitch while Δ*pilU*^m^ can. This agrees with recently established picture in literature that in *P. aeruginosa* PilT is the main contractile protein while PilU acts as a supporting PilT-dependent retraction motor (22, 26, 27). We note that this conclusion may not be true in other species, as in *D. nodosus* it has been shown that mutation of either *pilT* or *pilU* eliminated cell ability to twitch (24). Although Δ*pilU*^m^ of *P. aeruginosa* can twitch, Δ*pilU*^m^ cells exhibited slower and more directional-persistent surface motion compared to WT^m^.

Secondly, in terms of collective behavior, our results showed that PilU affects microcolony formation in two ways. On one hand, the loss of PilU would slow down the twitching movement of cells and temporally lead to a delayed microcolony formation. A similar observation of PilU loss in delaying the microcolony formation was also reported in *Neisseria meningitidis* (*N. meningitidis*) (40). However, unlike the hyperpiliation of Δ*pilU* in *P. aeruginosa*, in the study of *N. meningitidis* (40), the electron microscopy measurements showed no difference in the morphology or piliation levels between Δ*pilU* and WT strains. Although there are no twitching measurements in this study, the delaying of microcolony formation was also observed in liquid medium under which the twitching motility does not work. Together, it seems to suggest that there might be different causes for the delaying of microcolony formation observed in *P. aeruginosa* and *N. meningitidis*.

On the other hand, the loss of PilU also changed the TFP-based cell-cell collision responses from touch-turn dominance in WT to touch-upright dominance in Δ*pilU*, which then resulted in different morphologies of microcolonies. The altered cell-cell interactions in Δ*pilU* changed significantly the colony expansion pattern of bacteria on semi-solid agar surfaces, as illustrated in Figure 5, where Δ*pilU* cells formed a three-dimensional multilayerd “fence”-like edge that greatly reduced the expansion rate, whereas Δ*pilT*, Δ*pilA* and WT all have a single-layer of cells at the front of colony expansion. Similar differences in the colony edges were also found in a stab-inoculated sub-surface twitching assay, where *pilU* mutant was found to form a thick subsurface colony with a distinctive fringe whereas the *pilA* and *pilT* mutants formed colonies with smooth edges (24). These results suggest that the “fence”-like edge is likely due to the intrinsic properties of Δ*pilU* rather than substrate variations.

As Δ*pilU* cells are hyperpiliated and are also able to perform extension and retraction, we would like to know the passive role of TFP (i.e., physical filament itself) and the active role of TFP (i.e., through TFP extending and retracting activities) in these colony expansions. Δ*pilT* mutants are hyperpiliated but unable to retract TFP, so we can deduce the passive role of TFP from the measurements of Δ*pilT* which showed that TFP filaments can indeed slow down the colony expansion. This is consistent with the passive role of TFP found in swarming behavior of *P. aeruginosa* (39). The mechanism of the passive role of TFP presumably arises from the adhesive function of TFP, which have been shown to play roles in surface attachment as well as cell aggregations (1, 26, 34, 41). However, the passive role of TFP cannot explain the observation that the edge morphology of colony expansion of Δ*pilT* is more similar to that of both Δ*pilA* (no pili) and WT than that of Δ*pilU* (i.e., forming a monolayer of cells in Δ*pilT*, Δ*pilA* and WT, but not in Δ*pilU*), suggesting that the active role of TFP in Δ*pilU* is more important for the formation of multilayered “fence”-like edge. The increased touch-upright responses in Δ*pilU* can contribute to the formation of “fence”-like edge, as more upright cells would facilitate the formation of an “interception network” to prevents cells from passing through easily and thus more cells would accumulate near the edge, leading to a thick “fence”-like edge.

However, the mechanism underlying the increased touch-upright responses in Δ*pilU* is not clear. Our measurements of TFP distribution on cell surfaces showed that compared with WT^m^, Δ*pilU*^m^ displayed a higher percentage of two-poles events while Δ*pilT*^m^ showed a reduced percentage. These results seem in agreement with Chiang’s observations that PilT was found to be localized to both poles while PilU was localized at the piliated pole (42). Thus, one possible scenario to increase the touch-upright responses might be that loss of PilU would facilitate more cells having TFP located at both poles, which can create a tug-of-war situation when TFP at both poles retract simultaneously. Such a tug-of-war situation could be interrupted by cell collision, which leads to an imbalance in TFP retraction forces, causing cells to pivot into an upright position upon collision. We note that it is very likely that other factors would also be involved in TFP-polarization, such as PilG and PilH, which have been shown to control the polarization of *P. aeruginosa* during mechanotaxis (43). Future studies using high-resolution microscopy techniques to visualize TFP dynamics during cell-cell interactions could help elucidate the precise mechanism underlying this touch-upright phenotype.

In conclusion, our results expanded our current understanding of the role of PilU in *P. aeruginosa*. Beyond that PilU acts as a PilT-dependent retraction motor and is required for generating high retraction forces such as twitching in a typical agar stab assay (23, 44), we further demonstrated the roles of PilU in cell surface behaviors. We showed that loss of PilU reduced cell twitching activity, delayed microcolony formation, altered cell response behavior after collision and slowed down the colony expansion by forming a distinctive “fence”-like colony edge. While our study provides new insight into PilU’s roles beyond twitching, several questions remain. Future research should investigate the molecular mechanisms by which PilU influences TFP distribution and cell-cell interactions. Additionally, examining the impact of PilU on biofilm formation and virulence in animal models would further our understanding of its physiological importance.

## MATERIALS AND METHODS

### Bacterial strains and growth conditions

All strains and plasmids used in this study are listed in Table 1. *E. coli* DH5α strain was used as host for plasmid construction and amplification, see the literature (32). *P. aeruginosa* strains were cultured on Luria-Bertani (LB) agar plate at 37℃ for 12 h, and appropriate antibiotics (200 μg/mL carbenicillin) were added if necessary. By shaking in a FAB medium with 30 mM glutamic acid (Sigma-Aldrich) and selected concentrations of arabinose at 220 rpm and 37 ° C for about 6 hours to OD _600 nm_ ≈ 0.3. The culture was then diluted to OD _600 nm_ ≈ 0.01 in FAB medium containing 0.6 mM glutamic acid and arabinose for injection into the flow chamber. For strain PAO1P*_BAD_*-pilAS99C (WT^m^), Δ*pilU*P*_BAD_*-pilAS99C (Δ*pilU*^m^) and Δ*pilT*P*_BAD_*-pilAS99C (Δ*pilT*^m^) 0.2% arabinose (Sigma-Aldrich) was added to the medium to control the production of mutant PilA. A fully automatic inverted microscope is used to observe the movement of bacteria on glass surfaces.

**Table 1.** Strains used in this study.

| Strains | Characteristics | Source |
| --- | --- | --- |
| PAO1 wild type (WT) |  | Lab stock |
| $\Delta pilU$ | Cannot produce PilU | Lab stock |
| $\Delta pilT$ | Cannot produce PilT | Lab stock |
| $\Delta pilA$ | Cannot produce PilA | Lab stock |
| PAO1P<sub>BAD</sub>-pilAS99C<br>(WT<sup>m</sup>) | Pherd-20T-pilA in PAO1 | This study |
| $\Delta pilUP_{BAD}$-pilAS99C<br>($\Delta pilU^m$) | Pherd-20T-pilA in $\Delta pilU$ | This study |
| $\Delta pilTP_{BAD}$-pilAS99C<br>($\Delta pilT^m$) | Pherd-20T-pilA in $\Delta pilT$ | This study |
| WT-GFP | GFP plasmid introduction into PAO1 | This study |
| Plasmids |  |  |
| pHERD-20T | pUCP20T with araC-PBAD, <sup>AmpR</sup> | This study |
| Pherd-20T-pilA | pUCP20T, ParaBAD-pilA(S99C), <sup>AmpR</sup> | This study |

### Flow cell

Before inoculating bacteria into the flow tank, the FAB medium containing 0.6 mM glutamic acid was rinsed with a microinjection pump (Harvard Apparatus) at a flow rate of 30 mL/h for 5 min. The medium flow was then stopped and 1 mL of diluted bacterial culture (OD _600 nm_ ≈ 0.01) was injected directly into the channel of the flow pool using a 1 mL syringe equipped with a needle. After inoculation, an incubation period of 5 min was allowed to allow the cells to attach to the surface, followed by 5 min flushing at a high flow rate of 30 mL/h to wash away the floating cells. After that, set the flow rate to 3 mL/h and started the image recording. In this work, the flow cell experiment was carried out at 30 °C.

### Pili fluorescent staining

Pilin labelling was achieved using Alexa Fluor^TM^ F488 C5-maleimide (ThermoFisher, A10254). Firstly, the injection pump was stopped, and 300 μL AF488-mal solution with a concentration of 0.5 mg/mL was injected into the chamber through the injection port, and immediately incubated in the dark for 15 min. The injection pump was turned on, and the fresh medium was rinsed at 30 mL/h for 5 min to remove the bound dye in the flow cell. The cell bodies and labeled TFP were then imaged using fluorescence microscopy on a Leica DMi8 microscope.

### Data collection

Single cell tracking: An EMCCD camera (Andor iXon) was used to capture images on a Leica DMi8 microscope equipped with a zero-drift autofocus system. The image size is 66.5 μm x 66.5 μm (1024 x 1024 pixels). These images were taken with a 100x oil lens (plus a 2x magnifying glass).

TFP phenotype and expansion rate data acquisition (fluorescence shooting): the images were recorded every 8 s. In order to reduce the influence of fluorescence irradiation on the physiological state of bacteria, the observation field of each sample was changed for 1 min of shooting, and the sample was changed for about 30 min of shooting. Twitching movement data acquisition: Leica software was used to integrate equipment to record bright field images every 3 s, and the total recording time was about 12 h. Group expansion tracking: The Nikon Ti2 inverted fluorescence microscope equipped with drift autofocus system and infrared camera Prime95B were used for bright-field and fluorescent dual-channel data acquisition. A 60x objective lens (plus 2x magnification) and NIS-Elements Viewer software was used. The bright-field images were recorded with one image per every 3s. The fluorescence images were recorded every 30 min, and the total recording time was about 12 h.

### Single cell tracking image analysis

In this study, bacterial tracking was performed in the same way as in previous studies (32, 41). Through bacterial tracking, a variety of bacterial information can be obtained, including cell centroid coordinates, cell orientations, cell instantaneous velocity, and mean square displacements (MSD) of cellsetc. Bacterial visit frequency analysis was done in the same way as in a previous study (36).

When comparing the motility data of WT^m^ and Δ*pilU*^m^ cells, it is necessary to first determine whether the cells are in a state of motion, and then perform statistical analysis on the motile cells of the two strains. The threshold for determining cell movement or non-movement was determined using the result of Δ*pilT* cells, as their speed are influenced by cell growth. Δ*pilT* cells have a measured speed distribution (from 100 cells) with an average speed of 0.41 ± 0.04 μm/min and a peak value of 0.32 μm/min, so a threshold of 0.45 μm/min was selected.

The determination of motile cells: A cell is defined as a motile cell if its movement speed is greater than 0.45 μm/min at any given moment during the time it appears in the field of view. If its speed remains below 0.45 μm/min throughout, it is defined as a non-motile cell.

To determine the fraction of movement time of cell, we first selected data within the first 3 hours after cell adhesion to the surface and before the formation of biofilm microcolonies. Then, a specific cell was chosen, and if its instantaneous movement speed exceeded 0.45 μm/min, it was classified as motile, otherwise, it was classified as non-motile. The ratio of the duration of cell movement to the total time the cell appears in the field of view is the proportion of time during which the cell is in motion. This ratio can be used to describe the level of cellular activity during the observation period. For pili length measurements, pili that had retracted when image recording began were excluded from the analysis. For the measurement of pili elongation (contraction) speed, only cells that performed a single pilin elongation (contraction) within a 1-minute time window were analyzed.

The determination of cell touch-turn or touch-upright: When the pili of a cell touch another cell (cell body or pili), the cell changes its orientation and movement direction, but in a plane parallel to the substrate, this phenomenon is defined as touch-turn. When the pili of a cell touch another cell (cell body or pili), the cell becomes in stand-up configuration, this phenomenon is defined as touch-upright.

### The colony expansion experiments

The colony expansion experiments were performed in a similar way as described in reference (45). In these experiments, 0.3% FAB agar-air interface was used to observe the expansion motion. The bacterial solution was cultured overnight to OD_600 nm_ ≈ 1, and 0.5μL was added to the center of the solidified medium, and then the bacterial solution was hung in the biosafety cabinet for 2 min, so that the surface water of the bacterial solution evaporated to facilitate observation; Put the device on a thin coverglass, and a 60x lens was used for observation. Images were recorded every 3 s. Fluorescence intensity of the near-boundary area of mixing region refers to the average fluorescence intensity of the area within a range of 100 pixels away from the boundary of the mixing region, while the leading front region fluorescence intensity refers to the average fluorescence intensity of the area of the leading front region.

Measuring expansion rate: The bacterial expansion rate was calculated as the ratio of the total area visited by the bacteria divided by the time period during which bacteria expanded. In order to identify edges, the Sobel operator was used to convolve the image, followed by median filtering. Then, set a threshold to segment the image and calculated the area covered by bacteria.

## ACKNOWLEDGEMENTS

The funders have no role in the study design, data collection and interpretation, or the decision to submit the work for publication. This work was supported by: National Natural Science Foundation of China 12374206 (KZ) National Key R&D Program of China 2023YFC3402401 (KZ) National Key R&D Program of China 2020YFA0906900 (FJ) China Postdoctoral Science Foundation 2023M740523(JZ) Sichuan Provincial People’s Hospital Research Fund of the Academy 2022B1013 (JZ)

## AUTHOR CONTRIBUTIONS

Conceptualization: KZ

Methodology: JZ, YL, YZ, SL, YS, FJ, KZ

Investigation: JZ, YL, YZ, SL

Visualization: JZ, YL

Supervision: KZ, FJ, YS

Funding acquisition: KZ, FJ, YS

Writing—original draft: JZ, YL, YS, FJ, KZ

Writing—review & editing: JZ, YL, YS, FJ, KZ

## COMPETING INTERESTS

The authors declare no competing interests.

## DATA AND MATERIALS AVAILABILITY

All data are available in the main text or the supplementary materials.

